# Biophysical Fluid Dynamics in a Petri Dish

**DOI:** 10.1101/2024.02.13.580063

**Authors:** George T. Fortune, Eric Lauga, Raymond E Goldstein

## Abstract

The humble Petri dish is perhaps the simplest setting in which to examine the locomotion of swimming organisms, particularly those whose body size is tens of microns to millimetres. The fluid layer in such a container has a bottom no-slip surface and a stress-free upper boundary. It is of fundamental interest to understand the flow fields produced by the elementary and composite singularities of Stokes flow in this geometry. Building on the few particular cases that have previously been considered in the literature, we study here the image systems for the primary singularities of Stokes flow subject to such boundary conditions —the stokeslet, rotlet, source, rotlet dipole, source dipole and stresslet —paying particular attention to the far-field behavior. In several key situations, the depth-averaged fluid flow is accurately captured by the solution of an associated Brinkman equation whose screening length is proportional to the depth of the fluid layer. The case of hydrodynamic bound states formed by spinning microswimmers near a no-slip surface, discovered first using the alga *Volvox*, is reconsidered in the geometry of a Petri dish, where the powerlaw attractive interaction between microswimmers acquires unusual exponentially screened oscillations.

## I. INTRODUCTION

Since its development in 1887 by the German physician Julius Petri [1] for the facilitation of cell culturing, extending the bacterial culture methods pioneered by his mentor Robert Koch [2], the Petri dish has become an integral part of any biology laboratory. While still primarily used for culturing cells, providing storage space whilst reducing the risk of contamination, its simplicity and functionality allows it to be used in a wide range of other contexts: in chemistry to dry out precipitates and evaporate solvents (e.g. when studying Liesegang rings [3, 4]) or in entomology where they are convenient enclosures to study the behaviour of insects and small animals [5, 6]. A Petri dish environment is also a simple and common setting in which to examine the locomotion of swimming organisms, particularly those whose body size is tens of microns to millimetres [7–11]. The boundary condition at the bottom surface of such a container can be approximated as no-slip, while the top of the fluid is stress-free. Hence, a general question is: how does confinement in a Petri dish alter the nature of the flow induced by motile organisms?

The framework to answer this question lies of course with Green’s functions. In low Reynolds number fluid mechanics governed by the Stokes equations [12], the most important such function corresponds to the flow induced by a point force in an unbounded fluid and decays as 1*/r*. First written down by Lorentz [13] and later denoted a Stokeslet [14], it has been used to solve a wide range of fluid dynamical problems (see Happel and Brenner [15] and Kim and Karrila [16] for general overviews). One powerful extension to the Stokeslet involves a multipole expansion similar to that in electrostatics. The fluid flow caused by the motion of an arbitrary rigid body through a viscous fluid can be represented as that from a collection of point forces at the surface of the body [16]. Expanding the Stokeslet produced at an arbitrary point on the body’s surface as a Taylor series about the center of the body and then summing these contributions in the far field, one obtains a perturbation expansion for the fluid flow induced by the body [17]. Regardless of the particular shape of the particle, the fluid velocity field will exhibit common features. The leading order 1*/r* term is still a Stokeslet, but at higher orders, one finds distinct singularities. In particular the 1*/r*^2^ term, denoted a force dipole, can be separated into a symmetric part, denoted a stresslet [18], that corresponds to a symmetric hydrodynamic stress applied locally to the fluid, and an anti-symmetric part, denoted a rotlet [19] (called a couplet by Batchelor [18])), corresponding to a local hydrodynamic torque that produces rotational motion.

A well chosen distribution of such Stokes singularities that exploits the inherent symmetries of the system in question can be used to solve Stokes equations in a wide range of geometries and biological contexts [16]. Figure 1 illustrates the breadth of this approach, giving examples of biological flows associated with each of the low order Stokes singularities. Although classically in biological fluid dynamics the stresslet is the most common Stokes singularity considered [20], one sees that all low order Stokes singularities arise in familiar contexts.

**FIG. 1.**
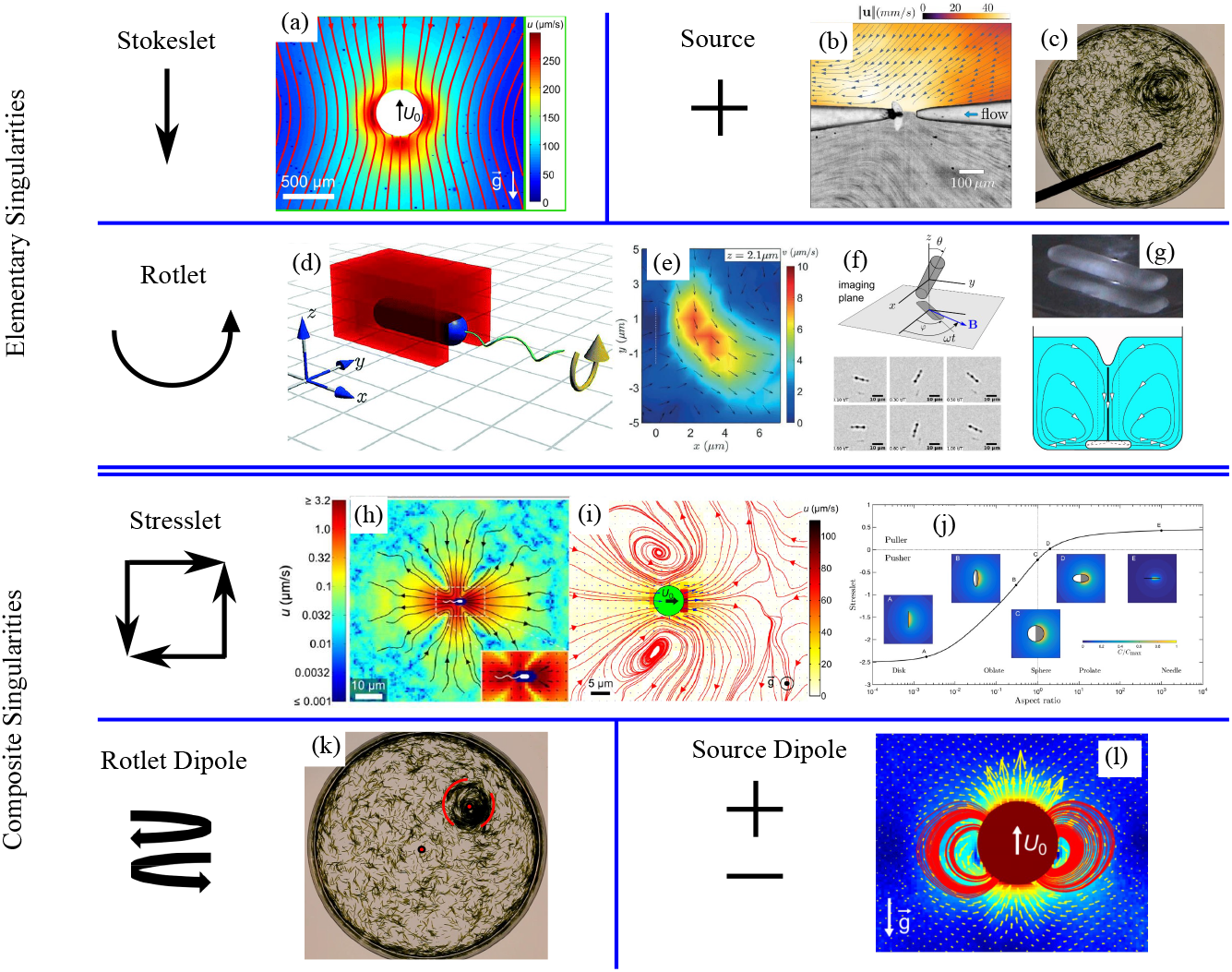
Stokes singularities in biological fluid mechanics. [a-g] Elementary singularities. Stokeslet flow is found in (a) far-field flow around *Volvox carteri* [21]. Source flows arise from injection of fluid from a micropipette into a Petri dish in studies of (b) dinoflagellates [22] and (c) plant-animal worms [23]. Rotlet flows arise from (d) the bacterium *Escherichia coli* under confinement, generating flow field in (e) [24], (f) a magnetic nano stir bar [25], and (g) a macroscopic stirrer [26]. [h-l] Composite singularities. Stresslets arise from (h) the pusher *E. coli* [9], (i) the puller alga *Chlamydomonas reinhardtii* [21], and (j) a phoretic Janus particle that changes from pusher to puller as a function of its aspect ratio [27]. A rotlet dipole flow is induced by (k) a circular mill of *Symsagittifera roscoffensis* [28]. A source is found in (l) the near-field flow induced by *Volvox carteri* after the Stokeslet contribution is subtracted [21].

The key question addressed here is thus: what is the fluid flow resulting from any Stokes singularity placed in a fluid layer between a rigid lower no-slip boundary and an upper stressfree surface. Although a few cases have been investigated in the literature, there has not been a systematic breakdown of the possible cases that arise. This was first considered by Liron and Mochon [29] who derived an exact solution in integral form for a Stokeslet. Subsequent work on this problem includes a theoretical study of bacterial swarms on agar [30], which contained a calculation of the leading order far field contribution to the flow from both a Stokeslet and a Rotlet when placed in a Petri dish configuration. This was further developed by Mathijssen, et. al. [31], who derived a numerically tractable approximation for the flow field produced by a Stokeslet and hence the flow field produced by a force- and torque-free micro-swimmer in a Petri dish.

In this paper, paying particular attention to the far-field behavior, we systematically extend and generalize these works beyond Stokeslets by computing exact expressions for the flow components *u*_*j*_ generated in a Petri dish of height *H* by the biologically relevant low-order primary and composite singularities of Stokes flow:

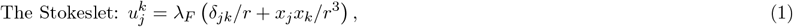

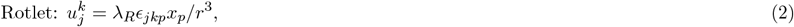

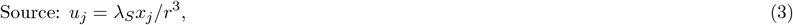

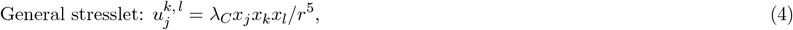

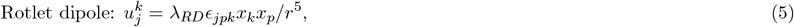

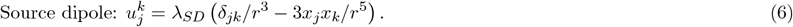

Note that here, *j, k* and *l* are free indices while the *λ*_*i*_ are dimensional constants denoting the strength of the singularities, with dimensions m^2^s^*−*1^ for the Stokeslet, m^3^s^*−*1^ for the rotlet, source and stresslet and m^4^s^*−*1^ for the rotlet dipole and source dipole. For clarity, we only present in the main text analysis for a source and a Stokeslet, namely the simplest and the most common singularity respectively. The results for the other singularities are given in Appendices B-E. Table I lists the locations of all these results in the paper. We adopt the geometry of Fig. 2, with in-plane coordinates (*x*_1_, *x*_2_), the no-slip surface at *x*_3_ = 0 and the stress-free surface at *x*_3_ = *H*.

**TABLE 1.**
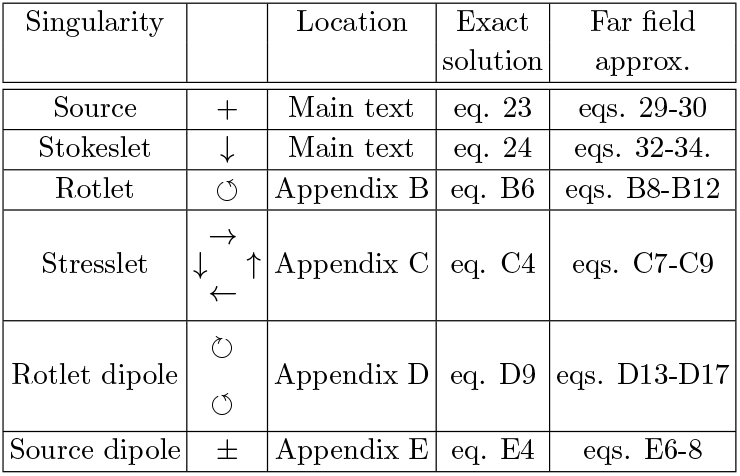
Location of results for various singularities.

**FIG. 2.**
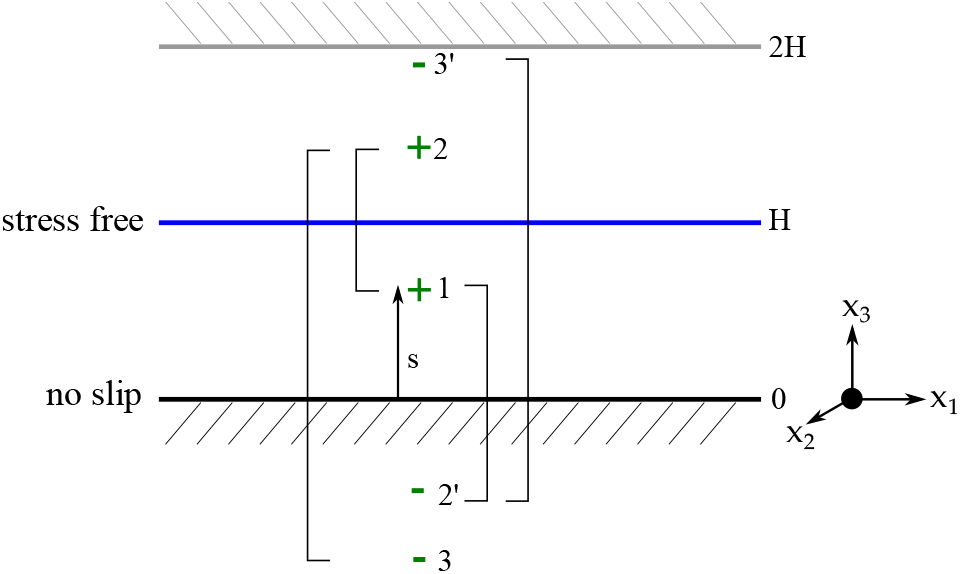
Stokes singularity in a Petri dish. The positive singularity is located at *z* = *s* and labelled 1. Its reflection across the no-stress surface at *z* = *H* is labelled 2 and across the no-slip surface at *z* = 0 is 2^*′*^, and so on. An alternate approach uses the full solution for a single no-slip surface and extends the domain to include a no-slip surface at *z* = 2*H*.

In §III, we calculate for both a source and a Stokeslet a particular solution to the Stokes equations generated by summing the infinite image system of Stokes singularities that is formed by repeatedly reflecting the initial singularity in both of the vertical boundaries. Then in §IV, an auxiliary solution is calculated using a Fourier transform method so that the sum of the two solutions is an exact solution for the full boundary conditions. In §V, a contour integral approach is used to calculate the leading order term of the fluid velocity in the far-field of a source.

This methodology, applied to both the source and the Stokeslet in §IV-V, is applied to the rest of the most commonly used Stokes singularities, (namely a rotlet, a general stresslet, a rotlet dipole and a source dipole), in Appendices B-E. Finally, as an application of these results, §VIII reconsiders in the geometry of the Petri dish the problem of hydrodynamic bound states, first discovered using the green alga *Volvox* near a no-slip surface [32] and later rediscovered in multiple contexts. The concluding §VI summarises the main results of the paper.

In particular, we note that higher order in-plane Stokes singularities can be found by differentiating the solutions with respect to a horizontal coordinate *x*_*α*_. Since all other Stokes singularities can be expressed in terms of derivatives of these singularities, we conclude that the leading order contribution to the fluid velocity in the far field for an arbitrary Stokes singularity is separable in *x*_3_, either decaying exponentially radially or having *x*_3_ dependence of the form *x*_3_(1*− x*_3_*/*2*H*). Hence, for many situations where the forcing can be modelled as a sum of Stokes singularities, the depth-averaged fluid flow can be captured by an associated Brinkman equation with a screening length proportional to *H*.

## II. SINGULARITY IN A PETRI DISH

Consider, as in Fig. 2, a Stokes singularity *f*, located at the point (*x*_1_, *x*_2_, *x*_3_) = (0, 0, *s*) between a rigid lower surface at *x*_3_ = 0 and an upper free surface at *x*_3_ = *H*, which generates a fluid flow ***u*** = (*u*_1_, *u*_2_, *u*_3_). At *x*_3_ = 0, we impose the no-slip boundary conditions

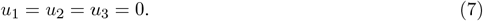

The capillary length *λ*_cap_ for a water-air interface is 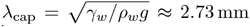, where *ρ*_*w*_ = 997 kgm^*−*3^ is the density of water, *γ*_*w*_ as 72.8 mNm^*−*1^ is the air-water surface tension, and *g* = 9.81 ms^*−*2^ is the gravitational acceleration. Since in a Petri dish *λ*_cap_ and *H* are similar in size, at the free surface, surface tension and gravitational effects are of similar magnitudes. Together, they restrict the vertical deformation of the interface. Hence, we assume the limit of no deformation in the vertical direction, fixing *H* as a constant. The self-consistency of this assumption is explored later in §VI. The dynamic boundary condition *u*_3_ = *DH/Dt* thus simplifies to

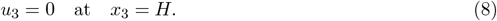

A force balance at 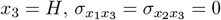, implies

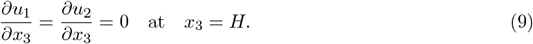

We nondimensionalize this system, scaling lengths with *H* and velocities with *U*_*S*_, where for a singularity of strength *λ*_*S*_ that decays in the far field like 1*/r*^*n*^, *U*_*S*_ = *λ*_*S*_*H*^*−n*^. For notational simplicity, we define

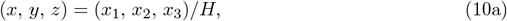

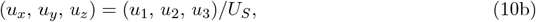

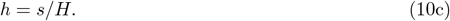

The boundary conditions become

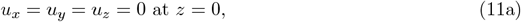

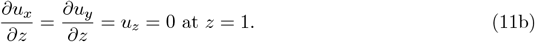

## III. REPEATED REFLECTION SOLUTION

We first examine the extent to which we can satisfy these boundary conditions through a distribution of image singularities. Following the canonical approach of Liron and Mochon [29], for a singularity placed at *x*_3_ = *s* [the green **+** labelled 1 in Fig. 2], placing an image singularity of the same sign at *x*_3_ = 2*H− s* (label 2) satisfies the free surface boundary condition at *x*_3_ = *H*. Similarly, placing an image singularity of the opposite sign at *x*_3_ = *−s* (2′^*′*^) partially satisfies the no-slip boundary condition, but singularity 2 fails the no-slip boundary condition and thus must be reflected about *x*_3_ = 0, changing its sign at location 3, Similarly, singularity 2^*′*^ fails the free surface boundary condition and thus must be reflected in *x*_3_ = *H* to give singularity 3^*′*^. Repeating this *ad infinitum*, namely inverting the sign when reflecting in the no-slip *x*_3_ = 0 boundary and keeping the same sign when reflecting in the free surface *x*_3_ = *H* boundary, gives an infinite series of singularities that constitutes the repeated reflection solution for that singularity.

In rescaled units, if we define the singularity locations ***r***_1*n*_ = (*x, y, z − h*+4*n*), ***r***_2*n*_ = (*x, y, z− h* + (4*n* + 2)), ***R***_1*n*_ = (*x, y, z* + *h* + 4*n*), and ***R***_2*n*_ = (*x, y, z* + *h* + (4*n* + 2)), then the repeated reflection solution is but one case of the general function *L*(*f*) for an arbitrary function *f*,

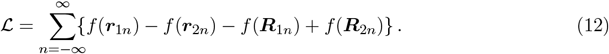

While intuitive, this series expansion is unwieldy. For the particular case *f* = 1*/r*, a Bessel function identity can be used to obtain the integral form

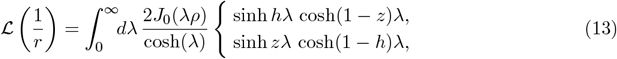

where 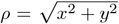 and here and below the upper expression holds for *z > h* and the lower for *z < h*. Higher order solutions are obtained from this result through algebraic manipulation, as shown in Appendix A for the third and fifth order cases. From those results, we find the repeated reflection solution *v*_*j*_ for a source *x*_*j*_*/r*^3^,

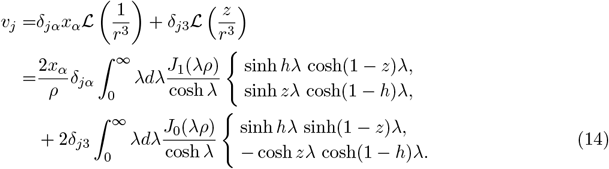

Similarly, for a Stokeslet δ_*jk*_*/r* + *x*_*j*_*x*_*k*_*/r*^3^, we find

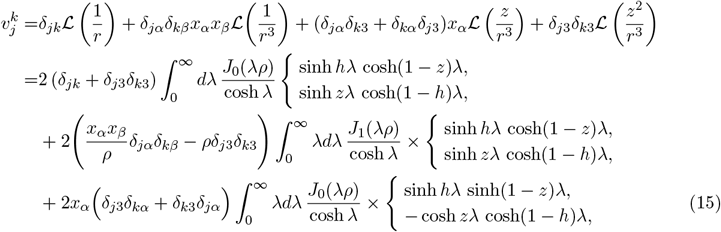

Similar expressions can be constructed for the other commonly used Stokes singularities (see Appendix B for the rotlet, C for the stresslet, D for the rotlet dipole, and E for the source dipole).

These results obtained via the repeated reflection solution can also be found directly from Liron’s solution [29] for a point force between two no-slip walls by setting the separation in that calculation to be 2*H*, placing a second force at 2*H− s* and observing that the reflection symmetry of the problem about the midline at *x*_3_ = *H* guarantees a stress-free condition at the midline.

Due to the nature of the algebraic manipulations performed above, these integral expressions do not converge when in the horizontal plane of the singularity *x*_3_ = *s*. Instead, it transpires that the correct integral expression to use instead is 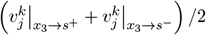, the average of the integrals as *x*_3_ tends to *s* from both directions.

## IV. AUXILIARY SOLUTION

In a scalar problem, such as a set of electric charges, the repeated reflection solution would solve the full system. However, our singularities are vectors and thus the repeated reflection solution does not satisfy all the boundary conditions. If we write the full fluid velocity field 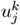 as 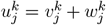, then the auxiliary solution 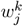 satisfies

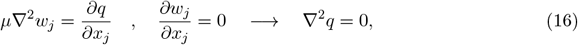

for suitable effective pressure *q*, with boundary conditions

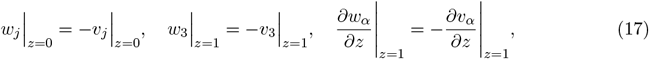

where *α* ∈ [1, 2] and *j* ∈ [1, 3]. For a source these are

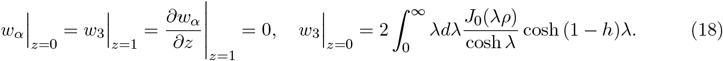

Similarly for a Stokeslet, applying standard Bessel function identities, the auxiliary boundary conditions become

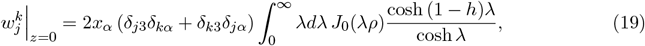

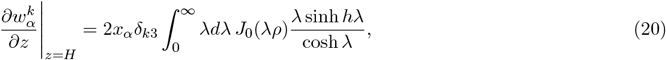

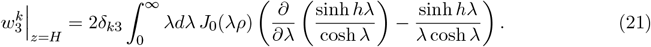

We solve for *w*_*j*_ by taking the two dimensional Fourier transform of this system with respect to (*x, y*), (namely *w*_*j*_(*x, y, z*) =⇒ ŵ_*j*_(*k*_1_, *k*_2_, *z*)), to arrive at

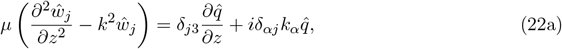

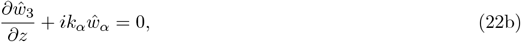

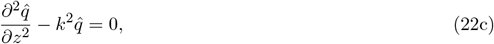

where *α* ∈ [1, 2] and 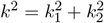. From inspection, this has the general solution

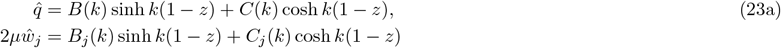

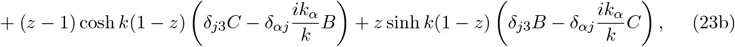

where {*B, C, B*_*j*_, *C*_*j*_}, with *j* ∈[1, 2, 3], are independent of *z*. From the continuity equation (22b) they satisfy

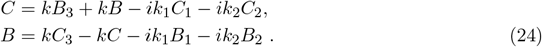

These constants are found on a case by case basis by transforming the boundary conditions given in (17) and solving through matrix methods the resulting set of eight coupled simultaneous equations in terms of {*k, h*}. For a source, (18) transforms to give

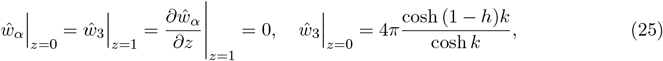

with corresponding full solution for *ŵ*_*j*_

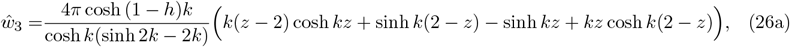

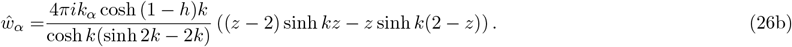

Similarly for a Stokeslet, (18)) transforms to give

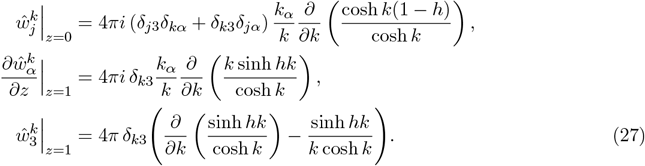

with corresponding full solution for 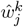

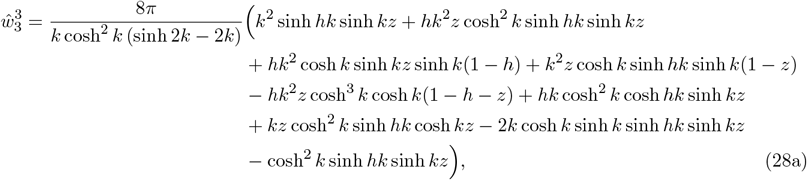

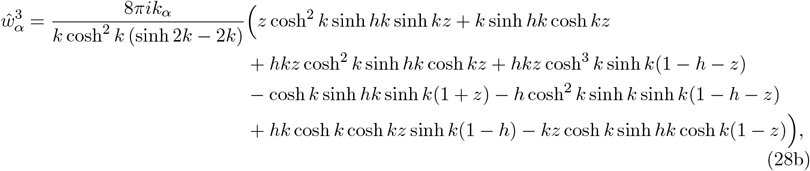

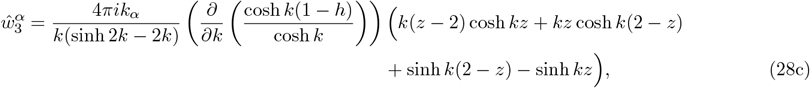

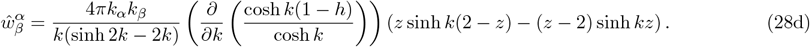

Rewriting the inverse Fourier transform in terms of Hankel transforms, we obtain for the source

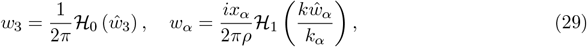

and for the Stokeslet

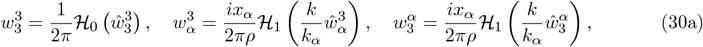

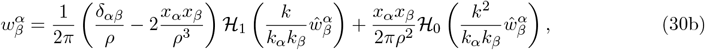

where *α*∈ [1, 2] and *ℌ*_*i*_ is the Hankel transform of order *i*. Similar integral expressions in terms of Hankel transforms can be constructed for other Stokes singularities (see Appendix B for the rotlet, C for the stresslet, D for the rotlet dipole, and E for the source dipole).

To illustrate the nature of these exact solutions, Fig. 3 plots various components of the fluid velocity field induced by four of the main singularities, the rotlet, source, rotlet dipole and source dipole, as a function of vertical height *z* for a range of horizontal radial distances away from the singularities, in each case located at *h* = 0.4.

**FIG. 3.**
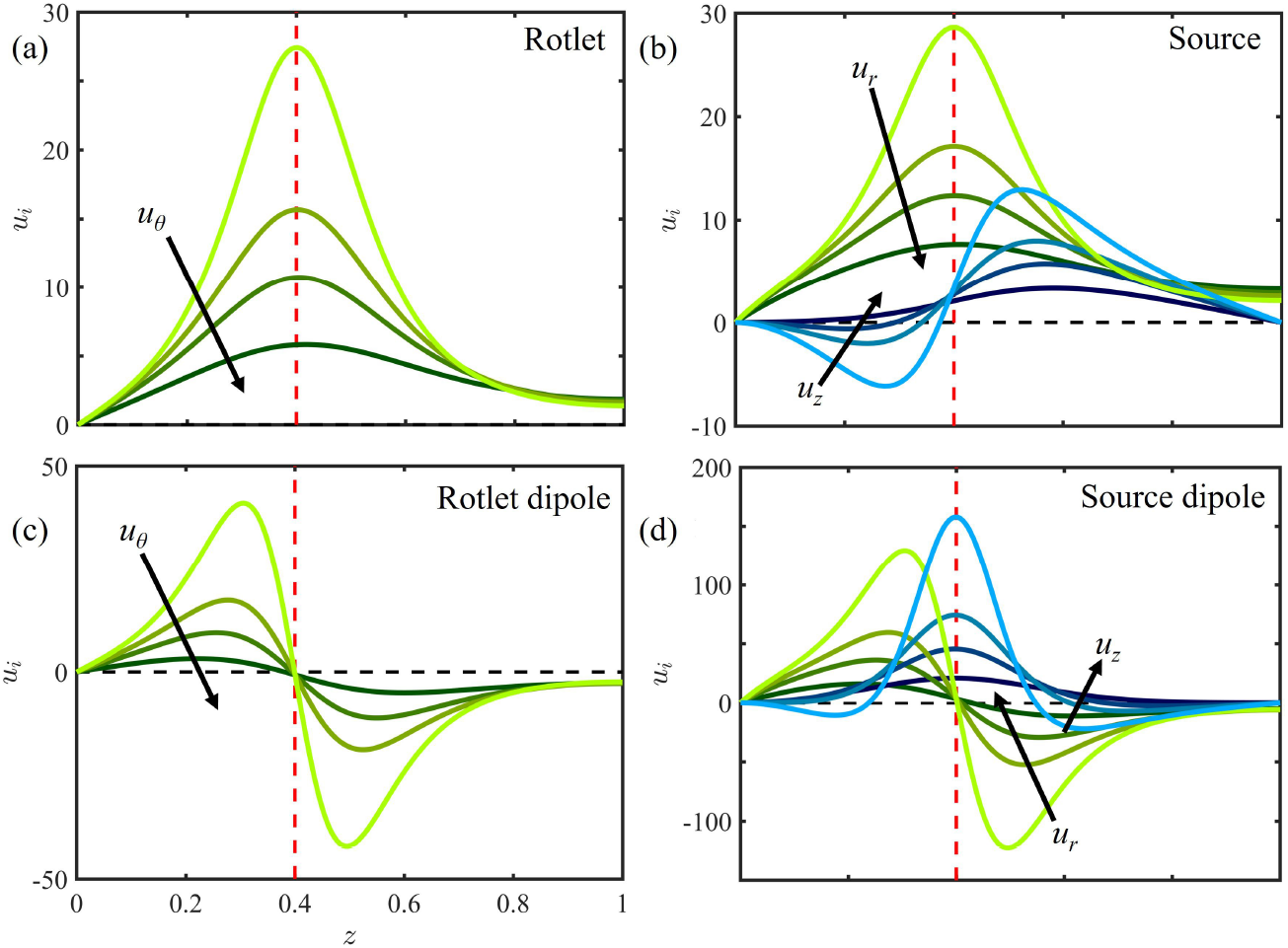
The near field velocity *u*_*i*_ produced by a number of singularities at *h* = 0.4 as a function of *z* for a range of *x* 0.19, 0.25, 0.3, 0.4, *y* = 0, with darker colours denoting larger *x*. (a) Rotlet, *i* = *θ* (green curves) (b) Source, *i* = *r* (green) or *i* = *z* (blue) (c) Rotlet dipole, *i* = *θ* (green) (d) Source dipole, *i* = *r* (green) or *i* = *z* (blue). Note that here (*r, θ*) are the polar coordinates for the horizontal plane i.e. *x* = *r* cos *θ* and *y* = *r* sin *θ*.

For the swirling component of the flow due to a rotlet, Fig. 3(a) illustrates clearly how the boundary conditions of no slip and no stress are satisfied, and the incipient divergence as the *x* location approaches that of the singularity. For the source in Fig. 3(b) the horizontal velocity *u*_*x*_ displays an increasing maximum as the observation point *x* approaches the singularity location, while the vertical velocity component *u*_*z*_ has a positive divergence for *z* → *h*^+^ and a negative divergence as *z*→ *h*^*−*^ as expected for a source, while vanishing at the top and bottom boundaries, as required by (11). Both the rotlet dipole in Fig. 3(c) and the source dipole in Fig. 3(d) appear as derivatives of their corresponding monopoles.

## V. FAR-FIELD SOLUTIONS

It is difficult to find the far-field (*ρ*≫ 1) behaviour of these solutions when they are expressed as exact solutions in integral form as Hankel transforms. Following the approach of Liron and Mochon [29], we may utilise a contour integration to express the exact solutions in series form. Given an even function *f* (*z*) decaying exponentially to zero on the real axis as *z* = *x* → *±*∞, consider the contour integral ∮_*γ*_ *F* where 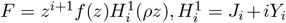 with *i* ∈ [0, 1] is a Hankel function of the 1st kind and *γ* = *γ*_0_ + *γ*_1_ + *γ*_*R*_ + *γ*_*ϵ*_ is a notched semicircular contour centered at the origin (Fig. 4). From Watson [38], 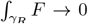 as *R* → ∞. Hence, applying the residue theorem in the limit as *R* → ∞ and *ϵ* → 0 yields

**FIG. 4.**
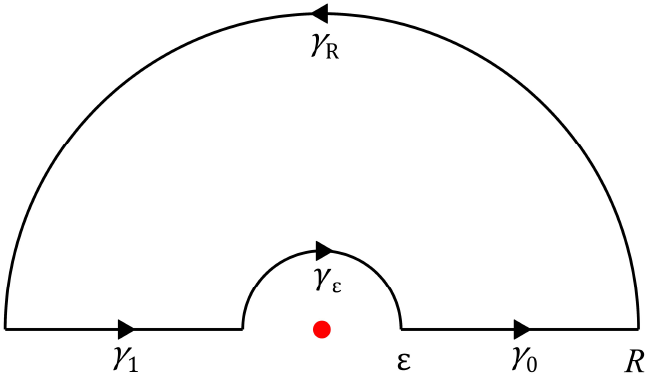
The notched semicircular contour *γ*.

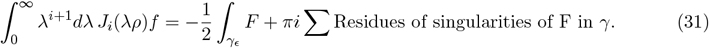

Using this method, the repeated reflection solutions *v*_*j*_ for all four primary Stokes singularities can be directly expressed in series form. For a source, *υ* _*j*_ becomes

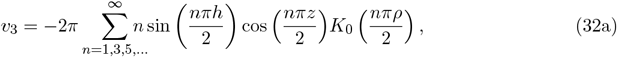

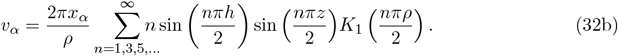

Note that for all four singularities, the dominant term in the far-field expansion (*ρ* ≫1) of the repeated reflection solution *v*_*j*_ comes from the *n* = 0 terms and decays like exp(*−πρ/*2). Similarly, the integral expressions for the auxiliary solution *w*_*j*_ can be expressed in series form to obtain series expansions for the full flow field *u*_*j*_. For a source, the corresponding complex function F has in *γ* poles of order 1 at *z*r= *πi*(*n* + 1*/*2) where *n* ∈ ℤ^≥^ and poles of order 1 at *z* = *z*_0_*/*2 where *z*_0_ satisfies sinh *z*_0_ = *z*_0_. Since 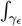 vanishes as *ϵ* → 0, when *j* = *k* = *l* = 3 (31) simplifies to become

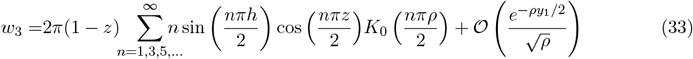

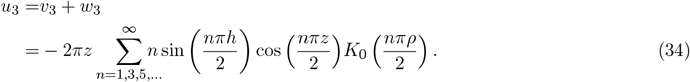

The first term dominates in the far-field, so

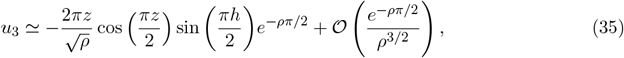

namely an exponential radial decay with *z* dependence *z* cos (*πz/*2), vanishing at both surfaces. Furthermore, when *j* = *α* ∈ [1, 2], the leading order contribution in the far field arises from *γ*_*ϵ*_, namely

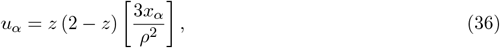

noting that the contribution from the poles at *z* = *πi*(*n* + 1*/*2) in *w*_*α*_ cancels out with *v*_*α*_. Similarly for a Stokeslet, *F* has poles of order 2 at *z* = *πi*(*n* + 1*/*2) where *n* ∈ ℤ^≥^ and poles of order 1 at *z* = *z*_0_*/*2 where *z*_0_ satisfies sinh *z*_0_ = *z*_0_. When *j* = *k* = 3, since 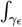 vanishes as *ϵ* → 0, (31) simplifies to

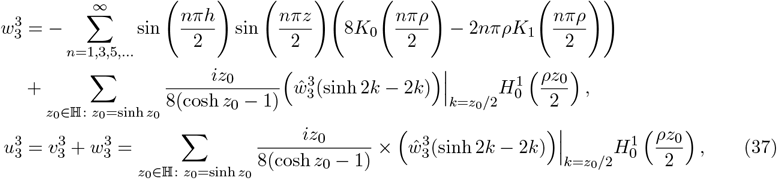

noting that the contribution from the poles of order 2 in 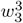 cancels out with 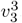. The leading far-field behavior is

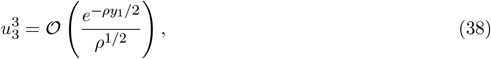

where *y*_1_ = 7.498 … is the imaginary part of the first non-zero root to sinh *z*_0_ = *z*_0_ in the first quadrant. Similarly for *j* = *α, k* = 3 and *k* = *α, j* = 3 where *α* ∈ [1, 2], the leading order far-field contribution is

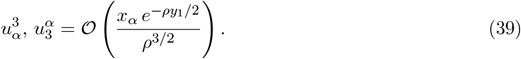

When *j* = *β* and *k* = *α* where *α, β* ∈ [1, 2], the leading order contribution in the far-field arises from *γ*_*ϵ*_,

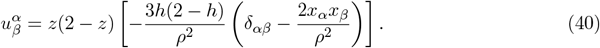

Similar far-field approximations can be found for the other Stokes singularities (Appendix B, rotlet; C, stresslet; D, rotlet dipole; E source dipole).

Figure. 5 plots streamlines of these far field flows in the horizontal plane *z* = 1. In Fig. 5(a), a Stokeslet orientated in the *x* direction generates a flow with a recirculating flow pattern of two loops decaying radially like 1*/ρ*^2^, namely a two dimensional source dipole (recalling that the source flow *u*_*s*_ = *x*_*i*_*/ρ*^2^ leads to the source dipole flow *u*_*sd*_ = δ_*ij*_*/ρ*^2^ *−* 2*x*_*i*_*x*_*j*_*/ρ*^4^). Confinement has fundamentally affected the unidirectionality of the flow by inducing recirculation in the *y* direction. This is a feature of the family of Stokes singularities that are derivatives of the Stokeslet, with higher order singularities having more recirculation loops. For example, a Stokes dipole has four loops while a Stokes quadrupole has six. In contrast, the spherical symmetry of a three dimensional source ensures that the new flow is still a source (Fig. 5(b)). Derivatives of the source, such as the source dipole, are also unchanged by confinement, and since the vertically orientated rotlet is independent of *z*, its streamlines are also unchanged, as seen in Fig. 5(c). Confinement breaks the symmetries of the horizontal rotlet and stresslet, leading to flows with the character a two dimensional source dipole for both a horizontally orientated rotlet (Fig. 5(d)) and a vertical stresslet (*j* = 1, *k* = 3, Fig. 5(e)) and a two dimensional source for a horizontal stresslet (*j* = *k* = 3, Fig. 5(f)), respectively.

**FIG. 5.**
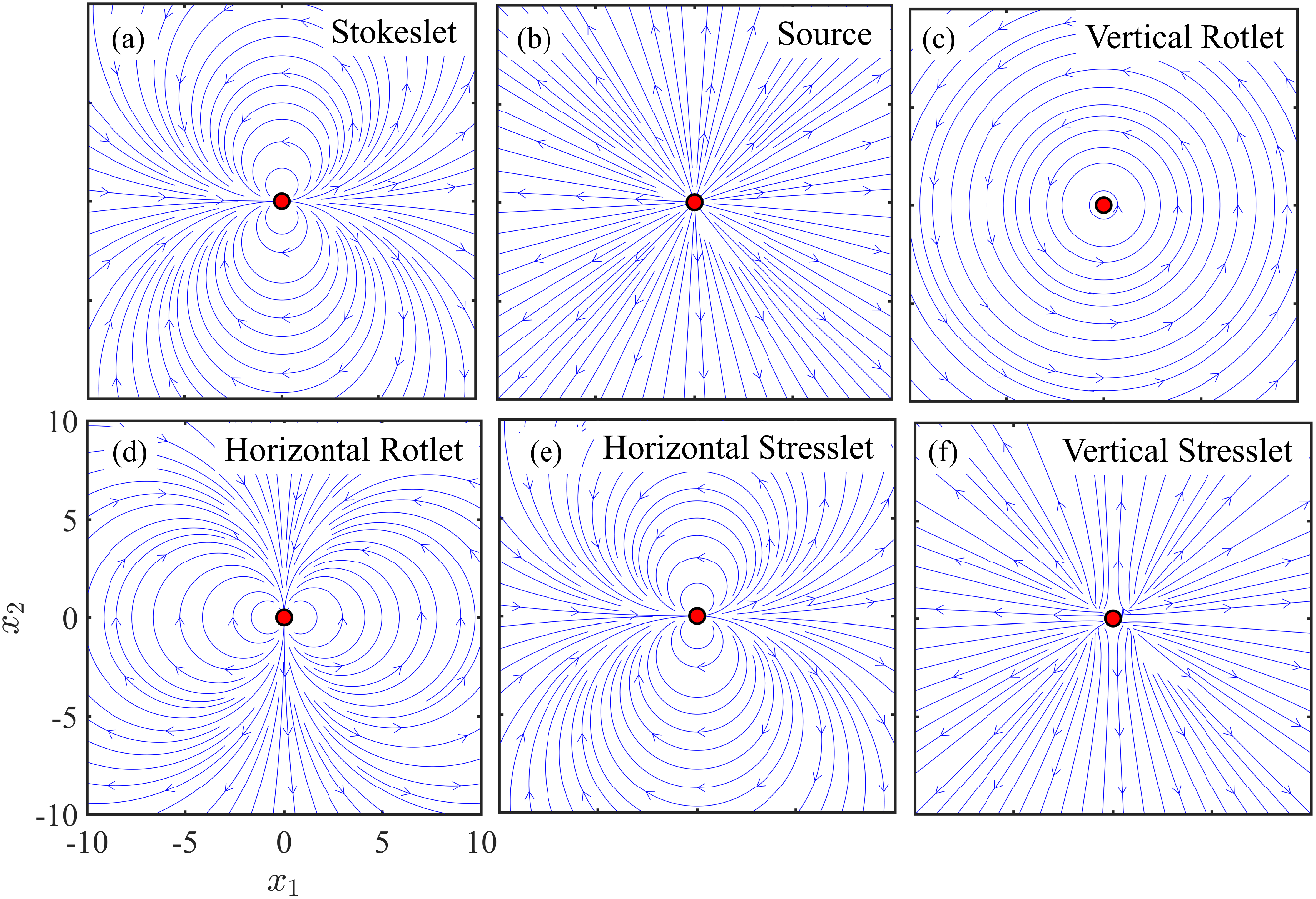
Streamlines in the *z* = 1 plane for the flows generated by Stokes singularities in the far field thin-film limit (*ρ* ≫*H*): (a) Stokeslet orientated in the positive *x* direction, (b) source, (c) and (d) Rotlet orientated in the *z* and *x* directions, respectively, (e) and (f) Stresslet *u*^*k,l*^ with *k* = 1, *l* = 3 and *k* = *l* = 1, respectively. As streamlines in (f) depend on *h*, we have set *h* = 1*/*2.

## VI. LEADING ORDER FAR FIELD FLOW

Examining the cases given above in §V and in Appendices (B)-(E), we note that for the four primary Stokes singularities, the leading order far-field flow is separable in *z* (formally considering the limit where *h, H, z* are fixed while *ρ* is large). If the flow does not decay exponentially radially, the it has *z* dependence of the form *z*(1 *− z/*2). Otherwise, the flow decays exponentially either as exp (*− ρπ/*2), arising from a *K*_1_(*− ρπ/*2) term with corresponding *z* dependence either sin *πz/*2 for horizontal flow or *z* cos *πz/*2 for vertical flow, or exp (*ρy*_1_*/*2) where *y*_1_≈ 7.498 is the imaginary part of the first non-zero root to sinh *z*_0_ = *z*_0_ in the upper half plane. All higher order Stokes singularities can be expressed as derivatives of these four primary Stokes singularities. These singularities must also either have leading order *z* dependence *z*(1 *− z/*2) or decay exponentially like exp (*− ρπ/*2) or exp (*− ρy*_1_*/*2). This means that the leading order far field contribution to the flow from these singularities can be obtained directly by differentiating the far field flows for the primary Stokes singularities, namely the full exact solutions which quickly become very complicated do not need to be derived. For example, differentiating (40) once, (40) twice and (36) once recovers the far field flows for a Stokes dipole, a Stokes quadrupole and a source dipole respectively given in [31], noting a sign error there in the expression given for a Stokes quadrupole (their equation (B8)), namely

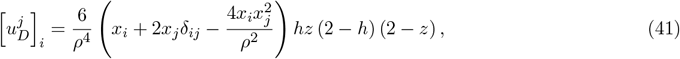

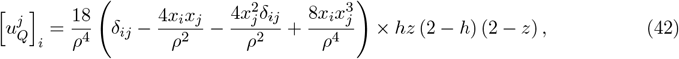

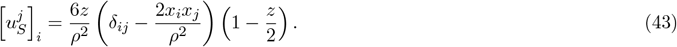

As a consistency check, (43) does indeed reproduce what was derived from first principles in Appendix E. Hence, for an arbitrary body whose free-space locomotion can be captured by a expansion in terms of Stokes singularities, the far field flow field is separable in *z* with either *z* dependence of the form *z*(1*− z/*2) or the flow decays radially exponentially. The fluid velocity field ***u*** can thus be factorised as ***u*** = *f* (*z*)***U*** (***x***_***h***_) where ***x***_***h***_ = (*x*_1_, *x*_2_) and *f* (*z*) is normalised so that 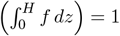 (typically f is either 3*z*(1 *− z/*2) or *π* sin (*πz/*2)*/*2). The 3D Stokes equations for ***u*** reduces to a Brinkman-like equation for the vertically averaged fluid velocity ***U***

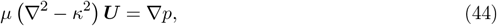

with corresponding incompressibility condition **∇ ·*U*** = 0, where *κ* = (∂*f/*∂*z*|_*z*=0_) ^½^plays the role of the inverse Debye screening length in screened electrostatics. We have thus reduced a 3D system to a 2D one that can be solved by transforming to an appropriate coordinate system that simplifies the boundary conditions. This method is equally applicable in the setup of Liron and Mochon [29], namely a microfluidic environment between two horizontal rigid boundaries, where the corresponding far field *z* dependence for an non radially exponentially decaying flow is *z*(1 *−z*).

## VII. SELF-CONSISTENCY CHECK

A key assumption made above was that the combination of surface tension and gravitational effects restricts vertical deformation of the interface and hence *H* can be assumed constant. As a self-consistency check, using (35), the leading order contribution to the stress 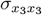 in the far field at the upper free surface boundary *x*_3_ = *H* that a source of strength *λ*_*S*_ (namely generating a flow *u*_*i*_ = *λ*_*S*_*x*_*i*_*/r*^3^) at (*x*_1_, *x*_2_, *x*_3_) = (0, 0, *s*) produces is

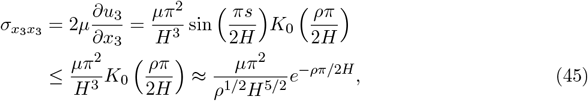

when *ρ* ≫ 2*H/π*. Here, we have utilized the asymptotic large argument expansion for *K*_*α*_ [39] together with the fact that | sin(*πs/*2*H*) | ≤1 ∀*s* ∈ [0, *H*]. Hence, a measure *M*_*s*_ of the relative strength of the stresses at the free surface arising from the flow generated by the singularity that seek to deform this surface to the gravitational forces restricting vertical deformation is

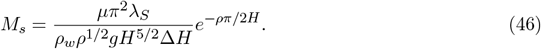

Writing the strength of the source *λ*_*S*_ as *λ*_*S*_ = *U*_*S*_*H*^2^, *U*_*s*_ scales with the typical velocities of flows in a Petri dish, namely *U*_*S*_ ∼ 2 mms^*−*1^. Hence, setting *µ* = 1 mPa s^*−*1^, *H* = 5 mm, Δ*H* = 0.1 mm, *ρ* = 1 cm we find *M*_*s*_ ≃1.2 ×10^*−*4^ ≪ 1, so *M*_*s*_ is indeed small and thus the flat surface approximation is consistent for a source.

## VIII. CASE STUDY: HYDRODYNAMIC BOUND STATES

An instructive application of the results of this paper is exploring the notion of “hydrodynamic bound states”. First discovered by Drescher, *et al*. in 2009 using the green alga *Volvox* [32], these are dynamical states exhibited by pairs of spherical chiral microswimmers near a surface. *Volvox* colonies have radius *R* ∼ 250 *µ*m, with ∼ 10^3^ biflagellated somatic cells beating on their surface. This beating is primarily in the posterior-anterior direction, but has a modest orthogonal component that leads to spinning motion about the AP axis. While the organisms are slightly denser than the fluid surrounding them, the flagellar beating allows them to swim upwards against gravity. When a suspension of *Volvox* was placed in a glass-topped chamber, the colonies naturally swam upwards due to their bottom-heaviness, which aligned their AP axis with gravity. Pairs of colonies at the chamber top were found to move together while they continued to spin, eventually touching and orbiting about each other.

As shown schematically in Fig. 6, once the colonies have ascended as high as possible, their centers are a distance *R* = *ϵH* (with *ϵ*≪ 1) below the upper no-slip surface. Due to their positive density offset relative to the surrounding ambient water, they are acted on by a downward gravitational force. Viewed from afar, each colony can be considered as a point force acting on a fluid: the resultant flow field is that of a downward-pointing Stokeslet of magnitude *F* = (4*π/*3)*R*^3^Δ*ρg* associated with the gravitational force. This geometry—two nearby Stokeslets directed away from a no-slip wall—is exactly that envisioned by Squires [33] in his analysis of surface-mediated interactions, who showed that the mutual advection of those Stokeslets toward each other is described by the dynamics of their separation *r* in the form

**FIG. 6.**
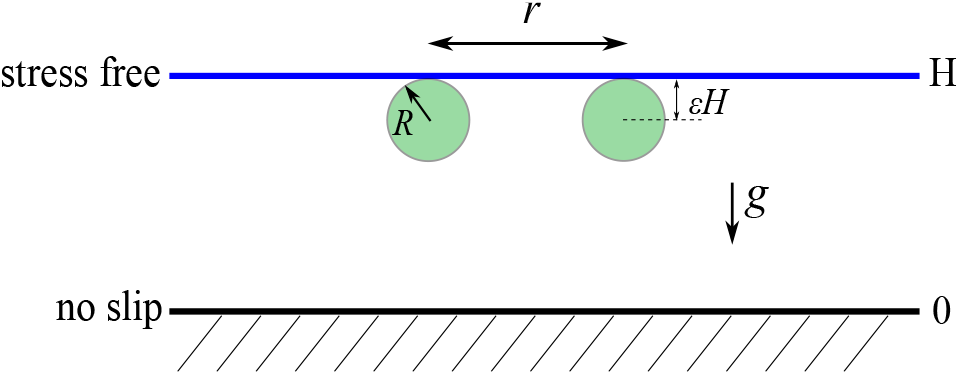
Geometry of hydrodynamic bound states. Two spherical, negatively buoyant microswimmers of radius *R* just below an upper surface, a horizontal distance *r* apart.

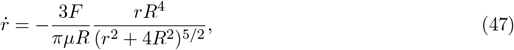

expressed in a way that identifies the characteristic speed *F/µR*. Tracking of *Volvox* pairs showed precise quantitative agreement with this result [32]. While it was not clear *a priori* that the Stokeslet approximation was valid over the large range of inter-colony separations explored, direct measurements of the flow fields around freely swimming colonies [21] showed that the Stokeslet does indeed dominate all higher-order singularities beyond a few radii from the colony center.

This general phenomenon has been rediscovered several times: in suspensions of the fast-moving bacterium *Thiovulum majus* [35], of the magnetotactic bacterium *Magnetotacticum magneticum* [36], and of starfish embryos [37]. In the latter case, the pairwise bound states occur at the air-water interface, which can be taken to be a stress-free boundary. In that case, and for an infinitely deep fluid, the image system for each Stokeslet is simply an opposite Stokeslet above the air-water interface - singularity 2 in Fig. 2. Thus, the lateral flow at (*x*_1_, 0, *x*_3_) due to a downward Stokeslet at the origin is

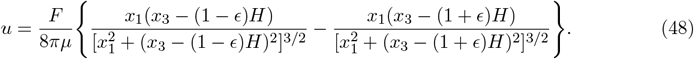

If we evaluate this flow at the Stokeslet location *x*_3_ = (1 *− ϵ*)*H*, and multiply by a factor of 2 we obtain the dynamics of the particle separation *r* in a form similar to the no-slip result (47), but with a different power law exponent in the denominator,

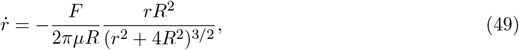

where *R* = *ϵH*. In each of (47) and (49) we can identify an effective potential energy *V* (*r*) such that 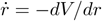. A natural question is how the result (49) for a stress-free surface is modified in the geometry of a Petri dish. The three lengths *R* = *ϵH, H*, and *r* must be compared to determine the appropriate asymptotic regime.

The dynamics (49) holds for *r*≪ *H* but without restriction on the relative sizes of *r* and *R*, except that the impenetrability of the colonies implies that this expression is only relevant for *r >* 2*R*. Of course, the validity of the singularity approach itself will decrease for *r* ∼ *R* = *ϵH*, and thus it is fair to assert that (49) is *physically* valid for *ϵH* ≪ *r* ≪ *H*, and in particular 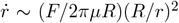 for *r* ≫ *R*. Indeed, as a consistency check, the full integral expressions do indeed simplify to (49) in this limit as we now show. Working in the same horizontal plane as the singularity, after some contour integration the repeated reflection solution becomes

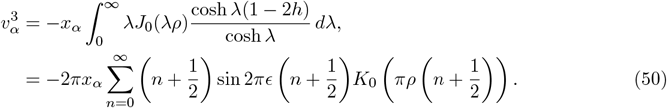

From (30a) we find that for small *ρ* the auxiliary solution 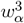 is 𝒪 (*x*_*α*_*ϵ/ρ*^2^) and hence, for points with small *ϵ* and *ρ*, the repeated reflection solution dominates the auxiliary solution. Expanding in powers of *ϵ* we find

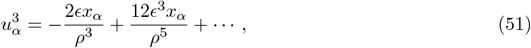

a result that agrees precisely with an expansion in *ϵ* and suitable nondimensionalization of (48).

The new regime of interest occurs when the separation *r* becomes comparable to or larger than the Petri dish depth *H*. Given for completeness in Appendix F, when *r* ≫ *H* (*ρ* ≫ 1), the non-dimensional flow field *u*^3^ decays exponentially with an unusual sinusoidal form

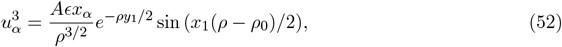

where *z*_1_ = *x*_1_ + *iy*_1_ = 2.769 + 7.498*i* is the first root in the first quadrant to the equation sinh *z*_1_ = *z*_1_, *A* = 38.340 and *ρ*_0_ = 0.298. Figure 7(a) explores this further, demonstrating how numerical solutions to the full flow field vary as a function of *ρ* for a range of values of *h*. Darker blue dots denote larger values of *h* i.e. the *Volvox* are closer to the free surface. For comparison, the asymptotic result (52) is superimposed on those numerical results. For clarity, all velocities are normalised by 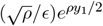 to highlight the sinusoidal component of the flow field. As can be seen, the asymptotic result is a good fit for *ρ* ≳ 2, improving as *ρ* increases and as *h* → 1.

**FIG. 7.**
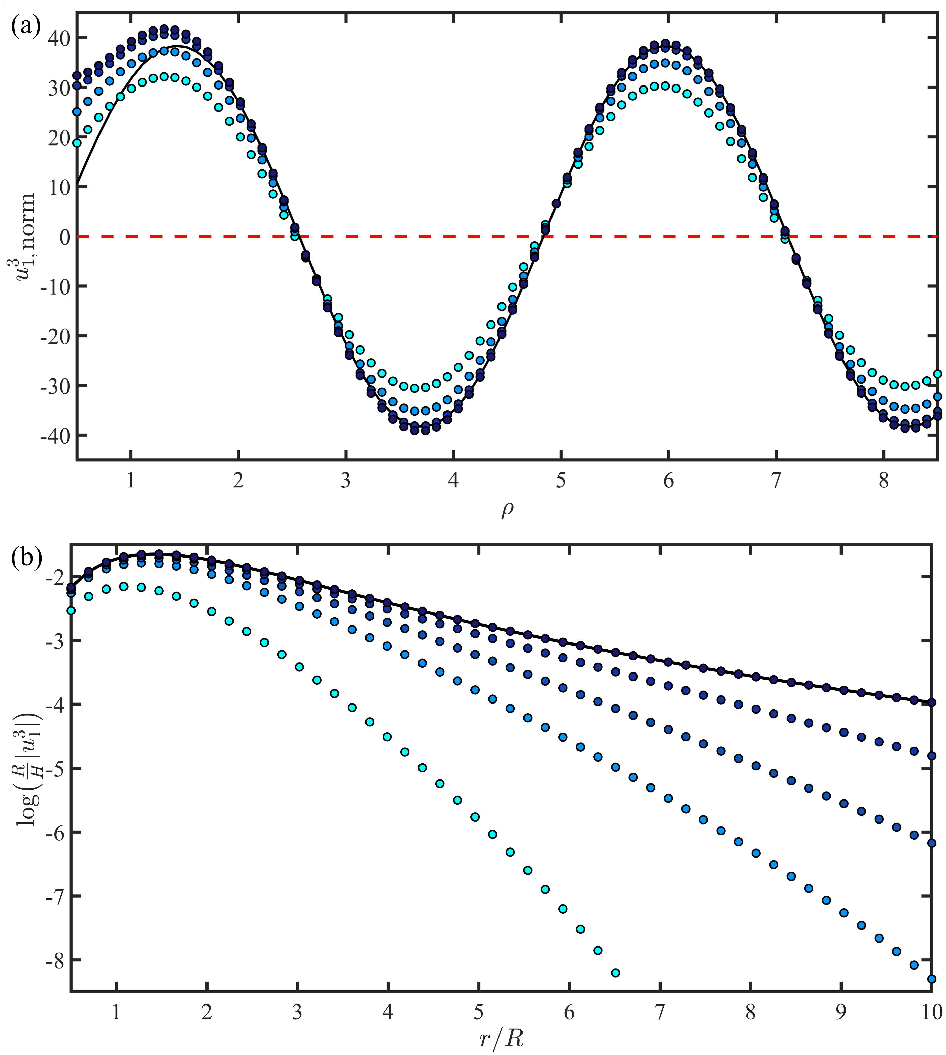
The lateral flow leading to hydrodynamic bound states. (a) Numerically obtained horizontal fluid velocity field 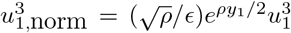, normalized to highlight the asymptotical sinusoidal component, generated by a vertically orientated Stokeslet placed at (0, 0, *H −R*) and evaluated as a function of *ρ* at the point (*ρ*, 0, *h*). Here, *R/H* ∈ [0.15, 0.1, 0.05, 0.01] with darker shades of blue denoting smaller values of *R/H*. The similarly scaled asymptotic result (52) is shown as the solid line. (b) The velocity *u*^3^ as a function of *r/R*. Here, *R/H* ∈ [0.3, 0.2, 0.15, 0.1, 0.01] with darker shades of blue denoting smaller values of *R/H*. For comparison the asymptotic result (48) for an infinitely deep Petri dish is shown as the solid black line.

An interesting feature of the screened interaction is that the multiplicative power law *ρ*^*−*1*/*2^ differs from that underlying the unscreened form (49), which falls off as *ρ*^*−*2^. This is unlike the case in electrostatics, for example, where a screened Coulomb interaction in three dimensions decays as ∼ (1*/r*)*e*^*−r/λ*^, where *λ* is the screening length, and the unscreened interation is ∼ 1*/r*. In the present case, the reason why we see a transition as *r* increases is that for small *r* the first reflection from the repeated reflection solution dominates, but as *r* increases the auxiliary solution generates terms that cancel out the repeated reflection solution, thus leaving lower order terms in the auxiliary solution to dominate, giving rise to an exponential decay.

Figure 7(b) shows in a semilogarithmic plot the lateral fluid velocity 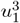 as a function of the dimensionless radial distance *r/R* for various values of *R/H*. The exponential cutoff of the powerlaw result (49) is evident. Even for the relatively large Petri dish depth *H/R* = 10 the velocity is attenuated by many orders of magnitude relative to the unscreened case for *r/R* ∼ 8, long before the sign oscillations are visible. Thus, while the corresponding evolution equation for the infalling of two colonies inherits the sign oscillations of the flow field (52), they appear only in the limit of very strong vertical confinement. The screening would, however, lead to very marked slowing down of the infalling trajectories relative to the infinite-depth case, and additionally reduce the significance of further-neighbor flows on a given swimmer in dense surface aggregates.

## IX. CONCLUSION

In this paper we have comprehensively explored the flows induced when Stokes singularities are placed in a Petri dish configuration, namely in a fluid layer with a bottom no-slip boundary and a top free surface boundary. In particular, we have derived both exact integral expression and far-field approximations for the flow generated by the four primary Stokes singularities: the Stokeslet, the rotlet, the source and the stresslet. Since all Stokes singularities can be expressed as derivatives of these four singularities, we can thus can gain insight about more general flows generated in a Petri dish by particles whose free space swimming fluid velocity can be represented as a sum of Stokes singularities. In particular, since the leading order contribution to the fluid velocity for these flows is separable in *z*, the full three dimensional Stokes equations can be vertically averaged to yield a much simpler two dimensional Brinkman equation much more amenable to analytic progress. A good example of this technique in action is [23], where the authors modeled a circular mill as a rotlet dipole, generating a radially exponentially decaying flow with *z* dependence sin (*πz/*2), and then solve the resulting Brinkman equation in the limit that the circular mill is away from the centre of the Petri dish by transforming to bipolar coordinates. We expect similar simplifications to hold in the many contexts in which experiments are carried out in the geometry of a Petri dish.

## ACKNOWLEDGMENTS

This work was supported in part by the Engineering and Physical Sciences Research Council, through a Doctoral Training Fellowship (GTF), by EPSRC grant EP/W024012/1 (EL,GTF), the European Research Council u nder the European Union’s Horizon 2020 Research and Innovation Programme (Grant No. 682754, EL), EPSRC Established Career Fellowship EP/M017982/1, Grant No. 7523 from the Marine Microbiology Initiative of the Gordon and Betty Moore Foundation, and the John Templeton Foundation (REG)

## Appendix A: Appendices Integral Notation and Higher Order Repeated Reflections Solutions

For clarity in Appendices B-E below, we define the functions *F*_*m, n*_ and *G*_*m, n*_

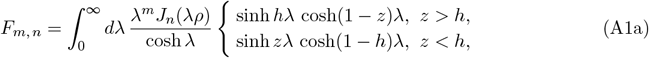

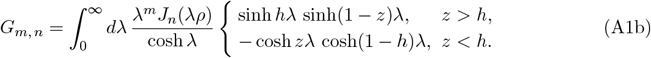

Shown in more detail elsewhere [28], these functions allow (13) in the main text to be extended to obtain repeated reflection solutions at third and fifth order,

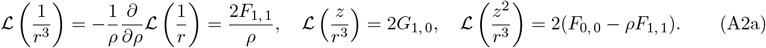

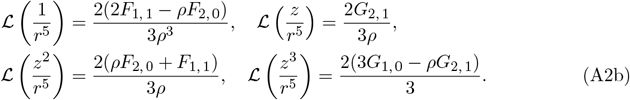

## Appendix B: Rotlet in a Petri Dish

The approach for a rotlet 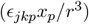 follows the procedure for the Stokeslet, with a repeated reflection solution

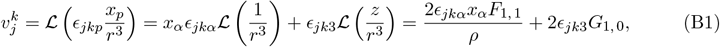

with the summation convention for *α* ∈ [1, 2]. The boundary conditions for the auxiliary solution 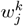 and transformed auxiliary solution 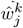 become

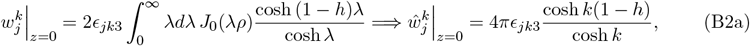

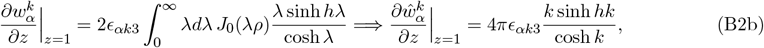

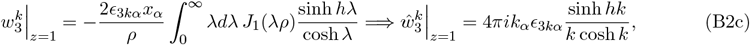

When k = 3 the boundary conditions are zero and 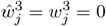. When *k* = *α* ∈ [1, 2], we find

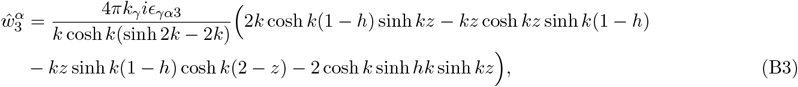

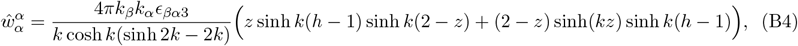

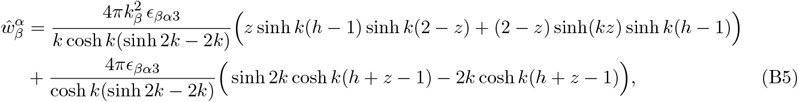

where *β* ∈ [1, 2] and *β* ≠ *α*. Rewriting inverse Fourier transforms in terms of Hankel transforms, we find

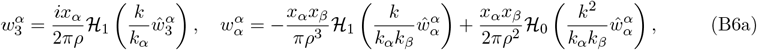

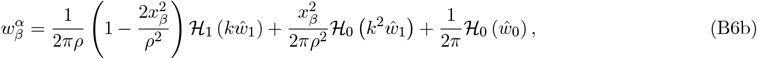

where *β* ∈ [1, 2], *β* ≠ *α* and for notational simplicity we have decomposed 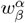 as 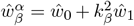. Using contour integration, as with the Stokeslet we find *F* has poles of order 1 at both *z* = *πi*(*n* + 1*/*2) (*n* ∈ ℤ^≥^) and *z* = *z*_0_*/*2 where *z*_0_ satisfies sinh *z*_0_ = *z*_0_. When 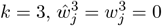, and thus the flow field 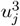 satisfies

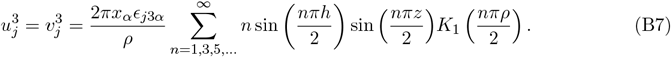

Hence in the far-field, the leading order contribution decays exponentially as

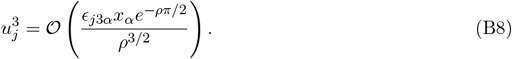

Since 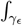 vanishes as *ϵ* → 0, when *j* = 3 and *k* = *α* where *α* ∈ [1, 2] (31) simplifies to

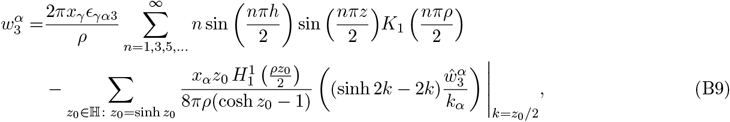

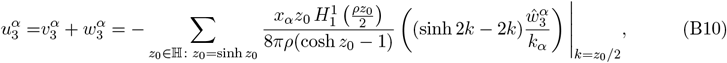

The contribution from poles of order 1 at *z* = *πi*(*n* + 1*/*2), *n* ∈ ℤ^≥^ cancels out with 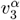, yielding

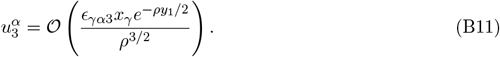

Finally, when *j, k* ∈ [1, 2], the leading order contribution in the far-field arises from *γ*_*ϵ*_ i.e.

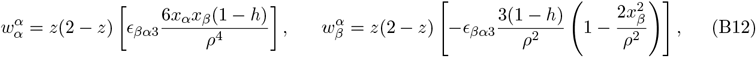

where *β* ∈ [1, 2] and *β* ≠ *α*.

## Appendix C: Stresslet in a Petri Dish

While the most general stresslet form is {*x*_*j*_*x*_*k*_*x*_*l*_*/r*^5^}, for swimming microorganisms typically *k* = *l*. From fifth order repeated reflection solutions, 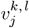 for a stresslet is

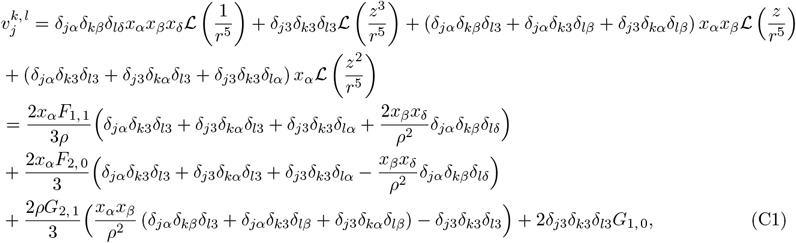

where {*α, β*, δ}∈ [1, 2]. The boundary conditions for the transformed auxiliary solution 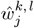 simplify to

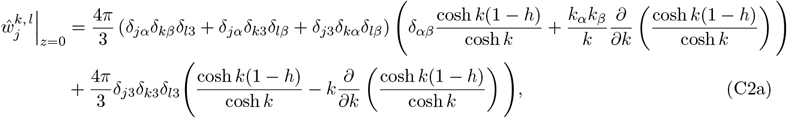

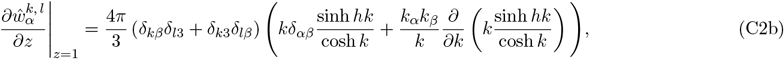

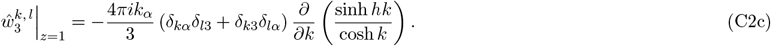

As in the main text for a source, we can thus solve for 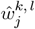 to give

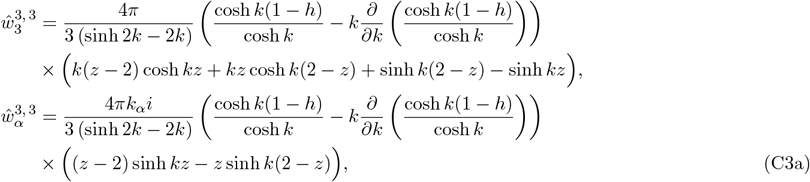

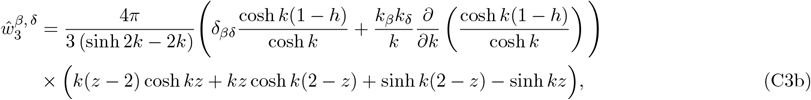

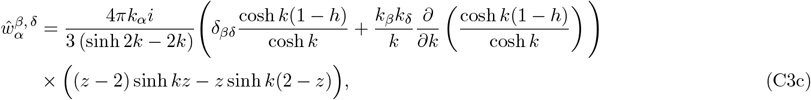

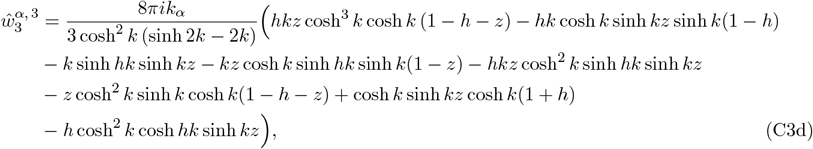

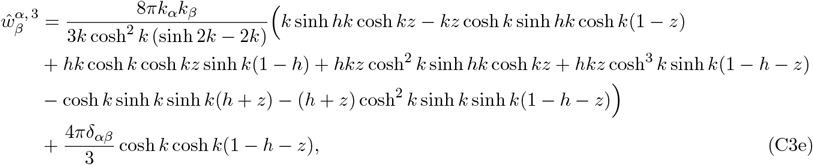

where *α, β*, δ ∈ [1, 2]. Hence, as above, we find the following integral expressions for 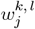.

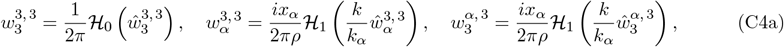

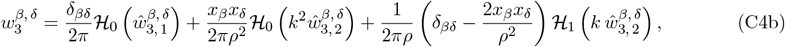

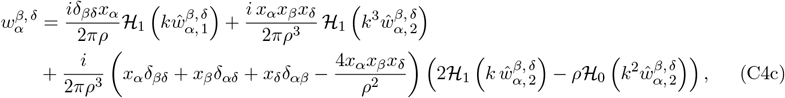

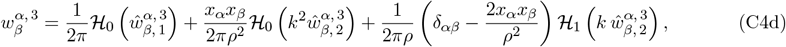

where for notational simplicity, we have decomposed 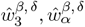 and 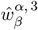as

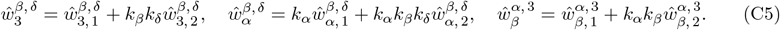

Similarly to the source above, F has in γ poles of order 2 at *z* = *πi* (*n* + 1/2) where *n* ∈ ℤ ≥ and poles of order 1 at *z* = *z*_0_*/*2 where *z*_0_ satisfies sinh *z*_0_ = *z*_0_. Since 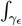 vanishes as *ϵ* → 0, when *j* = *k* = *l* = 3 (31) simplifies to become

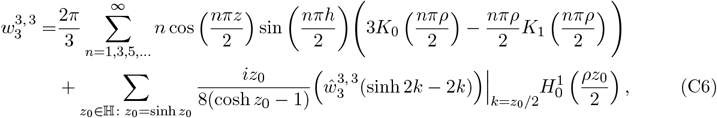

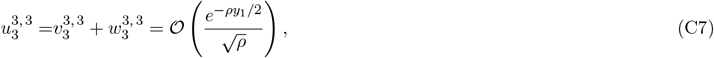

noting that as for the Stokeslet, the contribution from the poles of order 2 in 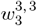 cancels out with 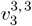. Similarly, the leading order contribution in the far-field when *j* = 3 for the other cases for *k* and *l* are

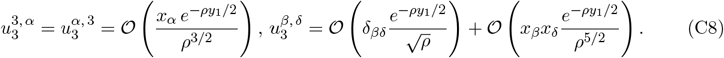

Finally, when *j* = *α* ∈ [1, 2], the leading order contribution in the far-field arises from *γ*_*ϵ*_, namely

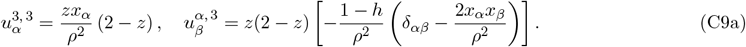

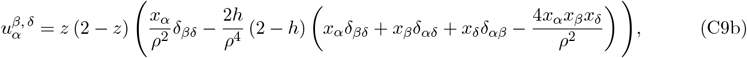

## Appendix D: Rotlet Dipole in a Petri Dish

From the fifth order repeated reflection solutions (Appendix A), 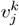 for a rotlet dipole is

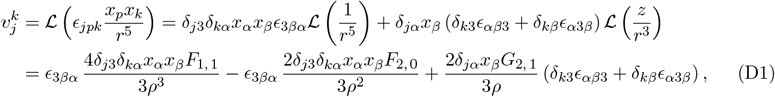

with boundary conditions for the corresponding auxiliary solution 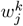 and transformed auxiliary solution 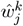

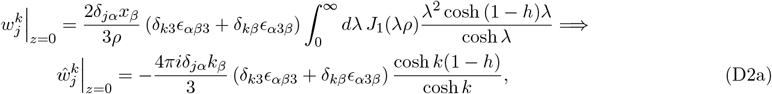

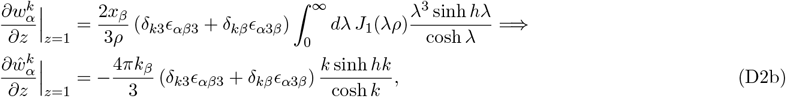

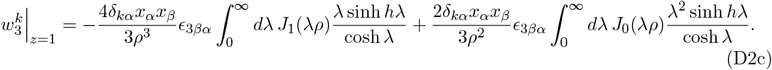

However, (D2c) is difficult to transform. Noting that *α* ≠*β* and utilising Bessel function identities, we find

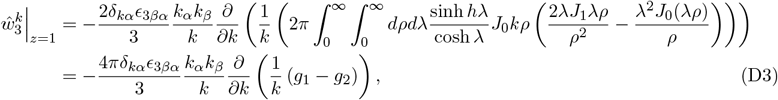

where *g*_1_ and *g*_2_ are defined as satisfying respectively

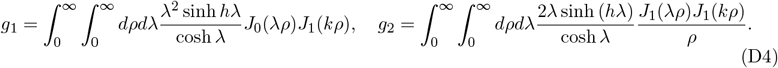

However, *g*_1_ simplifies to give

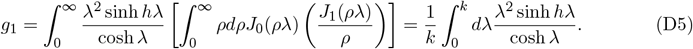

Furthermore, *g*_2_ simplifies to give

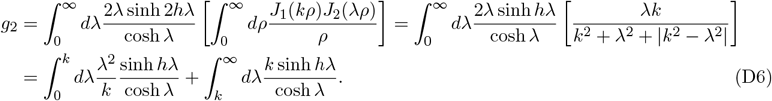

Putting this all together, (D3) becomes

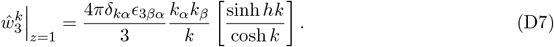

Hence, as in the main text for a source, we can thus solve for 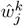 to give

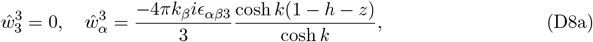

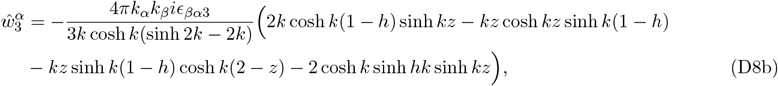

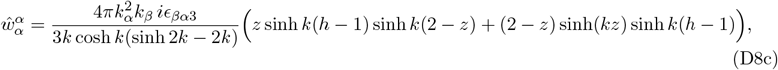

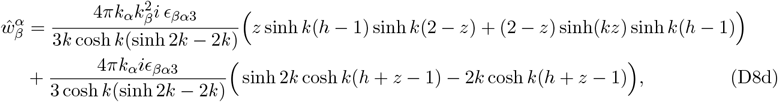

where *β* ∈ [1, 2] and *β* ≠*α*. Rewriting the inverse Fourier transform in terms of Hankel transforms, we get the following integral expressions for 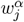

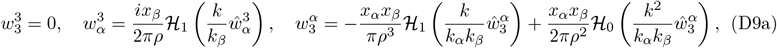

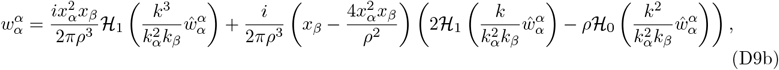

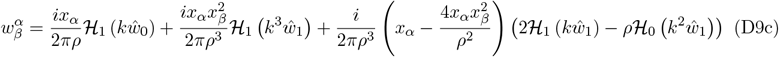

where *β* ∈ [1, 2], *β* ≠ *α* and for notational simplicity we have decomposed 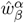 as 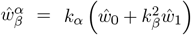. When *k* = *j* = 3, 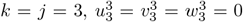. Furthermore, when *k* = 3 and *j* = *α*, we have

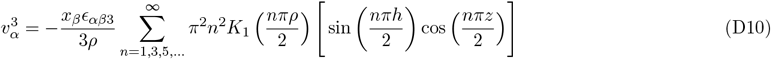

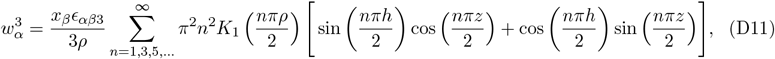

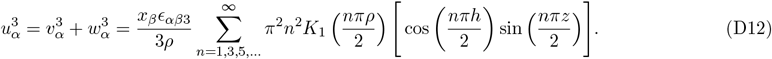

Hence in the far-field, the leading order contribution decays exponentially as

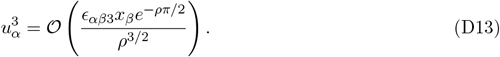

Since 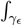 vanishes as *ϵ* → 0, when *j* = 3 and *k* = *α* where *α* ∈ [1, 2] (31) becomes

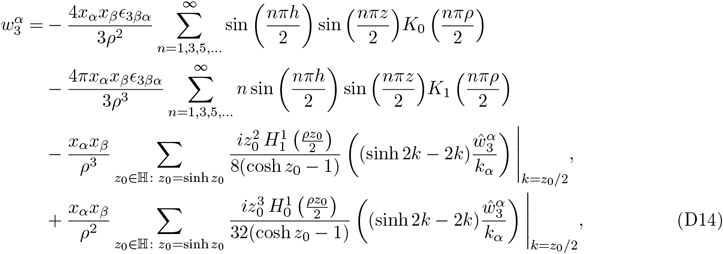

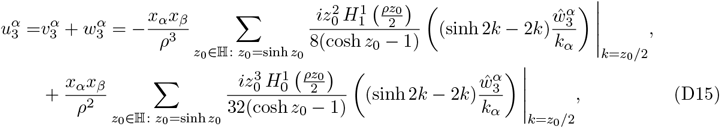

noting that the contribution from the poles of order 1 at *z* = *πi*(*n* + 1*/*2) where *n* ∈ ℤ^≥^ cancel out with 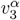. Hence,

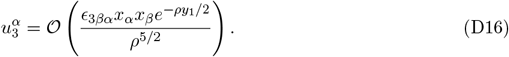

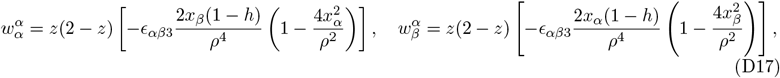

where *β* ∈ [1, 2] and *β* ≠ *α*.

## Appendix E: Source Dipole in a Petri Dish

Using fifth order repeated reflection solutions (Appendix A), 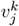for a source dipole becomes

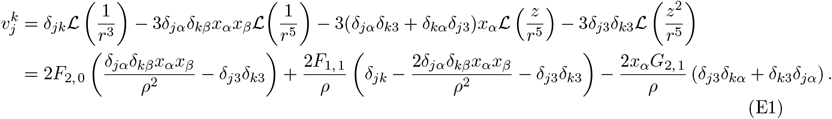

The boundary conditions for the corresponding auxiliary solution 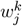 and transformed auxiliary solution 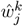 become

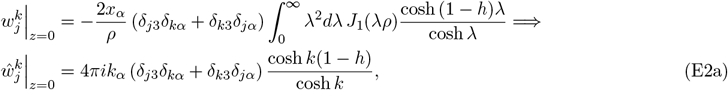

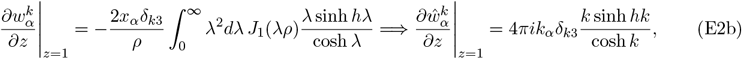

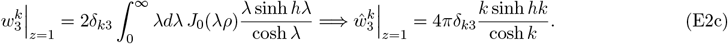

We thus obtain

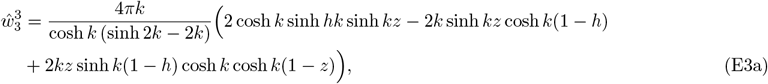

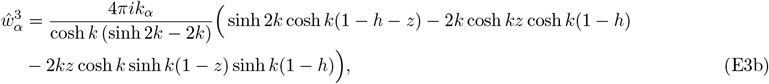

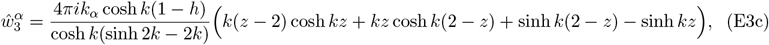

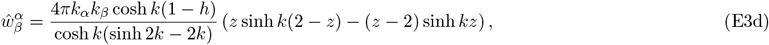

or, utilizing Hankel transforms,

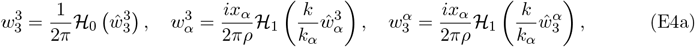

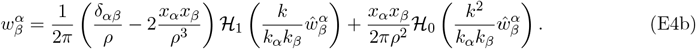

Similarly to the source, *F* has poles of order 1 at *z* = *πi*(*n*r+ 1*/*2) where *n* ∈ ℤ^≥^ and poles of order 1 at *z* = *z*_0_*/*2 where sinh *z*_0_ = *z*_0_. When *j* = *k* = 3, since 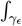 vanishes as *ϵ* → 0, and (31) becomes

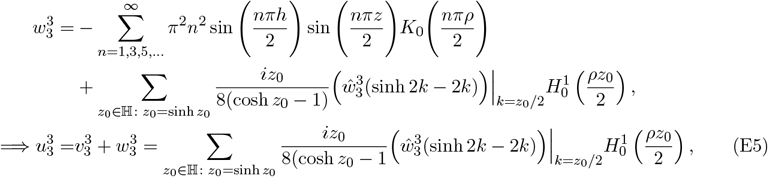

Hence, in the far-field the leading order contribution to 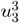 is

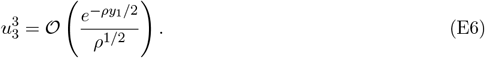

The leading order contribution in the far-field is

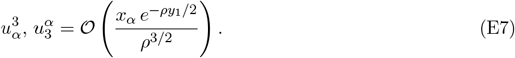

When *j* = *β* and *k* = *α* where *α, β* ∈ [1, 2], the leading far-field contribution arises from *γ*_*ϵ*_,

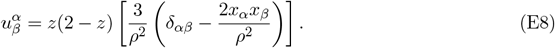

## Appendix F: Vertical Stokeslet Near the Free Surface Boundary

Here we find the leading term in (39) for the horizontal flow field at (*ρ*, 0, *h*) produced by a vertical Stokeslet located at (0, 0, *h*) in the limit that *ϵ* = 1 *− h* ≪ 1. From (28d) we have

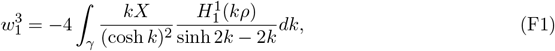

Where

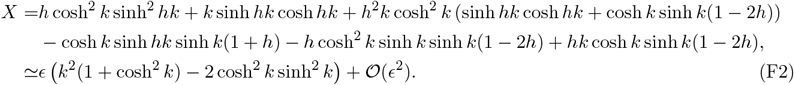

The leading order term of 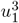 is that from 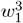 which is the sum of the residues at the first two roots in the upper half plane to the equation sinh 2*k* = 2*k* i.e. 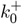 and 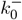 where

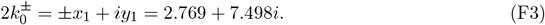

Hence, using the residue theorem we have

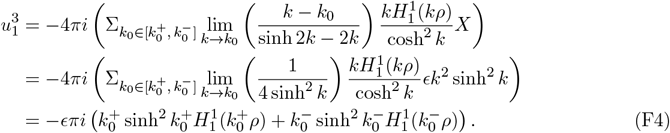

However, recall the standard result (e.g. see equation 9.2.3 of [39]) that

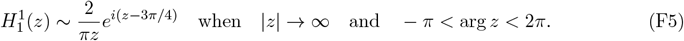

Hence, (F4) simplifies to become

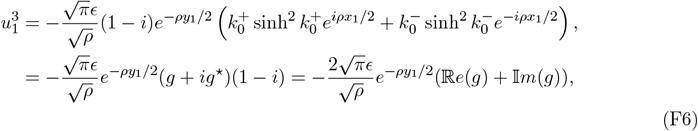

where *g* satisfies

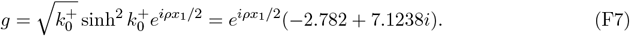

Rearranging this expression gives the relation in (52), namely

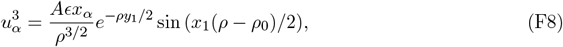

where *A* = 38.340 and *ρ*_0_ = 0.298.

